# Unravelling the glycome in human intervertebral disc degeneration: Aberrant glycosylation modulates inflammation and metabolism

**DOI:** 10.1101/2024.03.26.585840

**Authors:** Kieran Joyce, Aert F. Scheper, Aung Myat Phyo, Roisin O’Flaherty, Richard Drake, Aiden Devitt, Martina Marchetti-Deschmann, Radka Saldova, Abhay Pandit

## Abstract

Intervertebral disc (IVD) degeneration is one of the major contributing causes of low back pain (LBP), a common health issue that imposes a significant socio-economic burden on society. Previous work has demonstrated a dysregulated glycome in animal models of IVD degeneration; however, the role of glycosylation in pathogenesis is unknown. The objective of this study was to characterise altered glycan expression in IVD degeneration and elucidate the functional role of this response. Glycans in human healthy (n=6) and degenerated IVD (n=6) were examined through UPLC-MS and MALDI-IMS. These findings were correlated with proteomic analysis by LC-MS and functional *in vitro* studies using RNA sequencing. IVD degeneration was associated with a hypersialylated *N-*glycome, predominantly α-2,6 linked sialic acid. Confirming hypersialylation, we investigated sialylation’s functional role through mechanistic studies using a sialylation inhibitor (3Fax-peracetyl Neu5Ac). Sialylation inhibition *in vitro* modulated inflammatory and metabolic pathways, demonstrating a functional role for glycosylation in IVD degeneration.

**Brief summary:** IVD degeneration is associated with altered glycosylation, a potential target for new therapies.

## Introduction

Low back pain, a global health concern, imposes a significant socio-economic burden on society. A key contributor to this pain is intervertebral disc degeneration (IDD), a complex condition characterised by intricate alterations in the extracellular matrix, cell phenotype, and inflammatory responses. While tissue engineering solutions have been developed to address the underlying mechanisms of IDD and restore disc physiology, the role of glycosylation in this pathogenesis remains largely unexplored.

Glycosylation, the post-translational modification involving the addition and modification of carbohydrates to lipids and proteins by glycosyltransferases and glycosylhydrolases/glycosidases, generates a diverse range of glycoconjugates (1). Glycosylation expression provides insight into the temporal and spatial regulation of glycans in tissue inflammation and degeneration (1). These glycans are pivotal in protein folding, trafficking, receptor expression, activation, intracellular signalling, and immunomodulation (2).

Previous research has laid the foundation by investigating glycan expression at the histological level in embryological notochord and foetal intervertebral discs (3–5). However, advancements in glycoprotein extraction and analysis have allowed for a more precise exploration of glycosylation in connective tissues, such as cartilage (6–8). The glycosylation profile of IVD has been studied in murine (9), bovine (10) and ovine (11) models for injury and degeneration. These studies have briefly investigated the overall glycosylation motif expression using lectin arrays. Nevertheless, a significant gap exists in the field regarding the comprehensive analysis of the human glycome, encompassing *N*-glycans and other glycan species. Given the central role of glycans in metabolism, proliferation, and differentiation mechanisms, they may play a critical role in regenerative responses within the IVD, differentiating healthy discs from aged, degenerated ones.

To further unravel the role of glycans in degeneration, we turned our attention to sialylation, a type of glycosylation motif that plays a key role in cell-to-cell interactions, cell signalling and inflammation (12,13). Previous studies have established the co-localisation of sialic acid moieties at the level of chondrocytes in articular cartilage (8,14,15) and thereby recognised them as key components of the interactions of these cells with the extracellular matrix (ECM). Moreover, the role of dysregulation of sialic acids and sialic acid linkage on the cell surface, which is fundamental for cell-to-matrix interactions and mechanosensing, was characterised (15).

In this study, we set out two primary objectives: 1) to comprehensively characterise the *N*-glycome of the human disc with spatial and temporal resolution to investigate the dysregulation of glycans in degeneration. We sought to understand these changes in the context of the associated proteome and the overall glycome (including *O*-glycans, glycosphingolipids, etc.), combining dedicated *N*-glycan analysis and lectin microarray. Subsequently, upon confirming the presence of hypersialylation, we aimed to uncover the functional role of sialylation in degeneration through a mechanistic study using a small molecule inhibitor of sialylation (3Fax-peracetyl Neu5Ac; Neu5Ac-inhib). Our collective findings suggest that the glycosylation response, especially sialylation, plays a functional role in degeneration (12). Furthermore, sialylation inhibition can potentially reduce inflammation and oxidative stress, making it a promising adjunct to regenerative therapies. The experimental design for this study is outlined in Figure 1.

**Figure 1.**
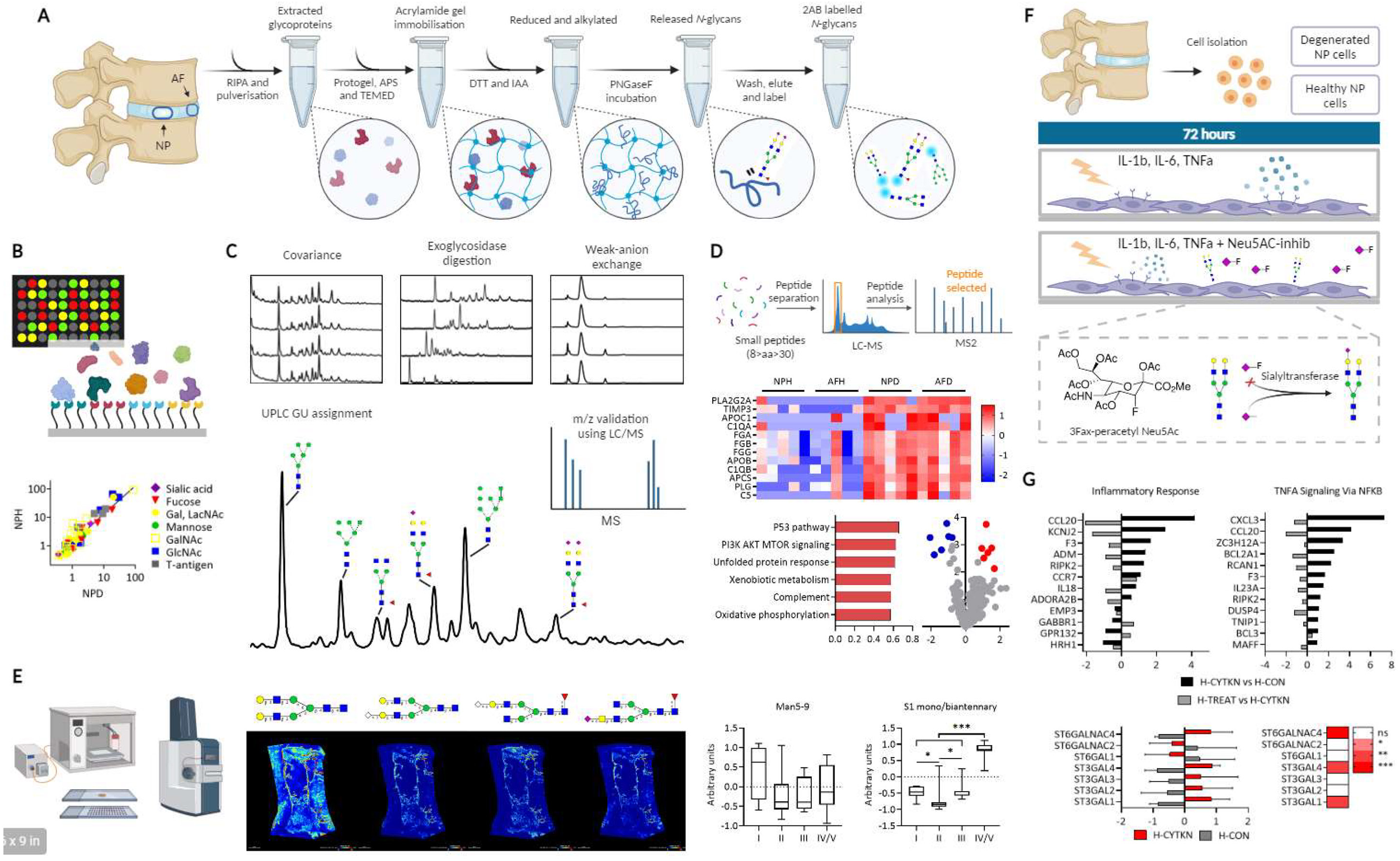
Schematic representation of the experimental design and procedures. A) Glycoprotein isolation and enzymatic *N*-glycan release. B) lectin microarray of intact glycoproteins. C) Quality assessment, assignment and quantification of *N*-glycans. D) Proteomic analysis workflow highlighting upregulated inflammatory pathways in the degenerated IVD. E) MALDI-TOF-MSI: glycan ion maps characterise the spatial presentation of *N*-glycans. F) Workflow for *in vitro* model of IVD degeneration and Neu5Ac-inhib studies. G) RNA seq data demonstrating upregulated inflammatory pathways and sialyltransferases under cytokine stimulation.

## Results

### Altered Nucleus Pulposus Glycosylation Motif Expression in Degeneration

The glycome is dynamic and responsive, present on protein, lipid, RNA and carbohydrate biomolecules through various conjugations. To investigate the presentation of glycosylated motifs in the IVD, lectin microarray was used to characterise the glycome in the human nucleus pulposus (NP) and annulus fibrosus (AF) in IVD degeneration (Table S1). Strong binding was observed for a wide range of lectin targets for all samples, indicating abundant glycosylated motifs (Figure 2A-B). Total lectin binding was greatest in the health NP (NPH), indicating that this tissue was most highly glycosylated, yet sample glycosylation signatures are conserved across this sample set. Overall, principal component analysis revealed the greatest separation of NPH from all other samples, with healthy AF (AFH) separated from degenerated tissues (NPD and AFD) by PC2 (Figure 2C-F). The glycomic profile of healthy NP differed from AF, containing significantly more mannose, galactose, T-antigen and *N*-acetylgalactosamine motifs (p<0.05, Fig 2G-M). Galactose, T-antigen and *N*-acetylgalactosamine motifs were all decreased in the degenerated NP (p <0.05), and there was no significant difference in degenerated NP and AF, indicating the NP expresses a more AF-like glyco-phenotype in degeneration.

**Figure 2.**
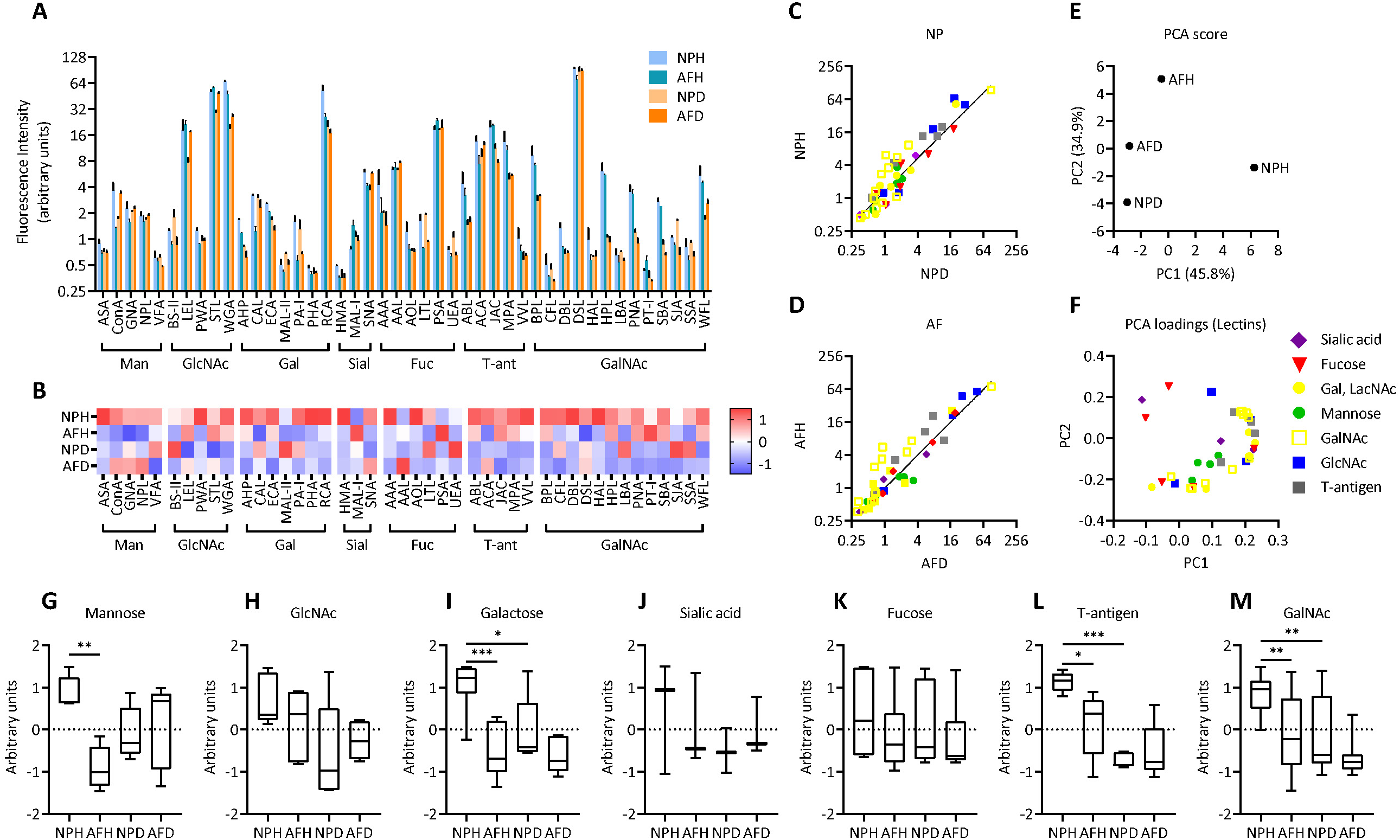
Glycosylation is downregulated in IVD degeneration. A) Fluorescence intensity of lectin binding in healthy NP (NPH), healthy AF (AFH), degenerated NP (NPD), and degenerated AF (AFH). B) Heatmap of z-score normalised lectin expression. C+D) Scatterplot of fluorescent intensities values for NP (C) and AF (D. E+F) Principal component analysis of lectin fluorescent intensities for each group (E) and loadings demonstrating individual loading contribution (F). G-M) Box plots of cumulative glycosylation motif expression represented as combined z-score normalised lectin expression. ANOVA was performed, followed by Tukey’s post hoc comparison. *p <0.05, **p <0.01, ***p <0.001.

### *N*-glycome of Human IVD

To unveil key glycans dysregulated in degeneration, the *N*-glycome from pooled healthy (Pfirrmann Grade I, n=6) and degenerated (Pfirmann Grade IV/V, n=6) IVD samples was released using the previously described method [22]. Six biological replicates from healthy and degenerated AF and NP were run undigested, where biological covariance varied from 5% to 70% (average below 20%) (Table S2). 32 peaks were identified as common across all individual sample replicates (Figure 2). The peaks with over 20% covariance accounted for less than 5% of *N*-glycans, occurring at the beginning or end of the profile with very low-intensity signal, where the small variability accounts for a greater proportion of peak variability increasing the covariance. Higher peak covariance was observed in degenerated samples, indicating heterogeneity across the glyco-signature in disease. Technical replicates demonstrated <5% covariance to ensure reproducibility as previously described (1).

Several techniques were employed to structurally assign peaks with glycan species. Each profile was digested by *Arthrobacter ureafaciens* sialidase (ABS) and run to confirm cleavable sialic acid content and highlight the presence of acetylation or other sialic acid modifications that ABS does not digest. Pooled profiles were also digested by a range of exoglycosidase enzymes to determine the structure and isomer of glycans in each assigned peak. 283 unique glycan species were identified with 270 species common across all *N*-glycan populations (Table S2). These were assigned to 48 glycan peaks common to the pooled samples with a summary of all assigned glycans and relative percentage abundance based on exoglycosidase digestion profiles and confirmed by LC-MS (GP1 – GP48; see Table S2). Significant differences in peak area were observed in GP5, 8, 10-15, 21 and 23 (Figure 3A-C). Glycan isoforms M5 (GP5), M8 (GP15) and FA3G3S2 (GP23) were significantly decreased in NPD vs NPH, while FA2G2 (GP10), FA2F1G1GalNAc1 (GP12) and FA2G2S1 (GP13) were significantly increased. FA2G2S2 (GP21) was the only peak significantly decreased in AFH vs NPH. M5 (GP5) was increased in AFH vs NPH while FA3G2 (GP11), FA2F1G1GalNAc1 (GP12) and FA2F2GalNAc2 (GP14) were all decreased. On the other hand, there were no significant differences in NPD and AFD, indicating that the tissue glyco-signature of the two tissues became similar, as observed in lectin microarray analysis.

**Figure 3.**
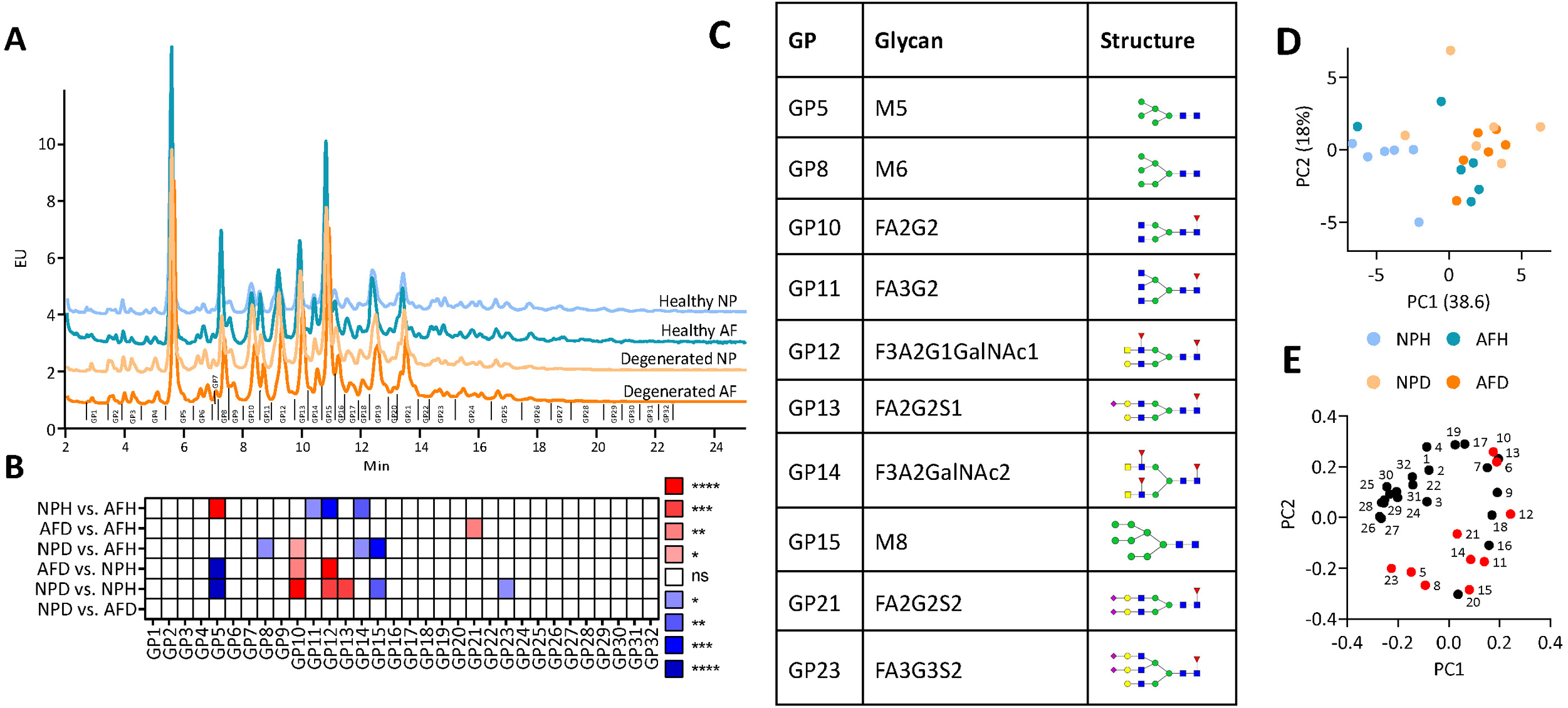
Bi-antennary and oligomannose glycans are differentially regulated in IVD degeneration, demonstrated using HILIC-UPLC. A) HILIC-UPLC chromatograms for *N*-glycans isolated from human IVD samples. 32 glycan peaks (GP) were common across all individual samples. The detailed composition of each peak is described in Supplementary Table S1. B+C) Summary of significantly altered peaks across group comparisons (B) and overview of the most abundant glycans in each significantly altered peak. Red indicates a significant increase and blue indicates a significant decrease in the relative area under the curve for each GP, *p <0.05, **p <0.01, ***p <0.001. (C). D+E) Principal component analysis and GP contributions to loadings. Red indicates significantly different GPs. Two-way ANOVA, Tukey’s post hoc test, p <0.05.

PCA was performed on the cumulative trends in glycosylation motifs (Fig 4A-B) and the complete set of 283 glycans, categorised by motif (Fig 4K-M), to gain a global overview of the glycomic data. The first two principal components described a combined variance of 81.6%, with PC1 and PC2 contributing 49.7% and 31.9%, respectively. Clear demarcation was observed between healthy NP, healthy AF, and degenerated tissues. PC1 provides a strong separation of healthy tissue, where oligomannose, tetraantennary, lactosamine and non-sialylated glycans are strong contributors towards a healthy phenotype, while biantennary, outer arm fucose, sialylated and complex glycans contributed to the degenerated phenotype. The amounts of total branching, sialylation, outer-arm fucose, oligomannose, LacNAc, GalNAc and substituent features of glycans were calculated based on the composition of these 48 peaks UPLC-MS analysis revealed specific trends in glycan species across the AF and NP in healthy and diseased tissues (Figure 4). Overall, monosialylated (S1) and biantennary (A2) glycans increased significantly in degenerated tissues vs healthy IVD (Figure 2D-E, Figure S1). Outer arm fucosylation trends towards an increase in degenerated NP vs healthy controls (Figure 2F). Conversely, oligomannose (Man5-9), tetraantennary (A4) and lactosamine terminated (LacNAc) glycans decreased in degenerated tissues, indicating a shift from branched high-molecular weight glycans to small monoantennary and biantennary glycan species (Figure 2G-J). Core fucosylation and galactosylation were not significantly altered. Sialylation linkages were characterised by differentiating between NAN1 and ABS exoglycosidase digestions to discriminate α-(2,3) vs α-(2,6) linkages. Sialylated α-(2,6) linkages were increased in degenerated IVD tissues from 19.57% and 19.46% in healthy AF and NP to 24.05% and 23.10% in degenerated AF and NP, respectively (Table S2).

**Figure 4.**
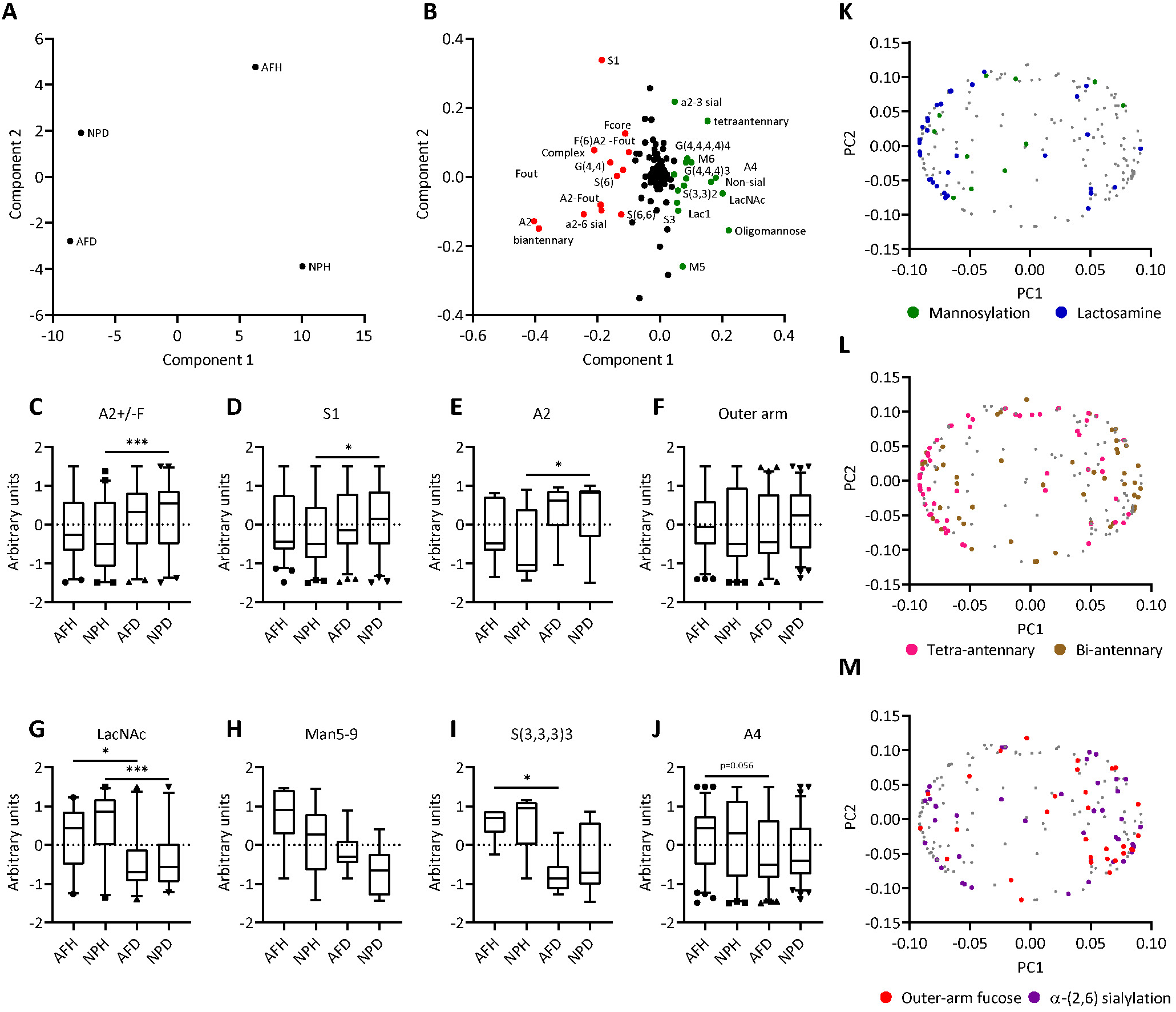
Differentially presented glycan traits in IVD degeneration. A+B) Principal component analysis and loadings demonstrating glycan trait contributions. C-J) Individual glycans were grouped into the following glycosylation traits; biantennary +/− outer arm fucosylation (A2+/−F) (C), monosialylated glycans (S1)(D), biantennary excluding outer arm fucosylated species (A2)(E), outer arm fucosylated (outer arm)(F), lactosaminylated (LavNAc)(G), higher oligomannose - from Man5 to Man9 (Man5-9)(H), trisialylated, alpha 2,3-linked (S(3,3,3)3)(I), tetraantennary (A4)(J). K-M) Principal component analysis loadings with highlighted traits: mannosylated (green) and lactoaminylated(blue) glycans (K), Tetraantennary (pink) and biantennary (brown) (L), and outer arm fucosylated (red) and alpha-2,6 sialylated (purple) (M). Two-way ANOVA, Tukey’s post hoc test, *p <0.05, ***p <0.001

### Spatial Glycan Analysis

MALDI-IMS of *N*-glycans was used to determine the spatial regulation of *N*-glycans in the human IVD, as described by Rebelo *et al.* (16). Human IVD tissue was antigen retrieved, sprayed with a molecular coating of PNGase F, and coated with CHCA matrix. This differs from UPLC analysis, where here only accessible *N*-glycans can be cleaved by PNGaseF. The enzymatically released *N*-glycans were detected with a MALDI–FTICR mass spectrometer. Glycan composition was confirmed by identifying the mass-to-charge (m/z) ratio of each peak and cross-referencing the Consortium for Functional Glycomics (accessible at www.functionalglycomics.org) database, within a PPM error of <20.0. This process was validated using the data previously obtained through HILIC-UPLC, allowing for enhanced refinement and a more comprehensive understanding of the *N*-glycome within the human IVD. An intra-grade comparison was carried out to build upon the prior analysis. Overall, 77 *N*-glycans were detected on MALDI MSI (Table S3). Distinct differences in glycan patterning were observed between grades of degeneration (Fig. 5). Oligomannose and tetraantennary *N*-glycan structures were more abundant in Grade I and II, while mono/biantennary, with or without sialylation, were more abundant in Grade IV/V (Fig, 5F-K).

**Figure 5.**
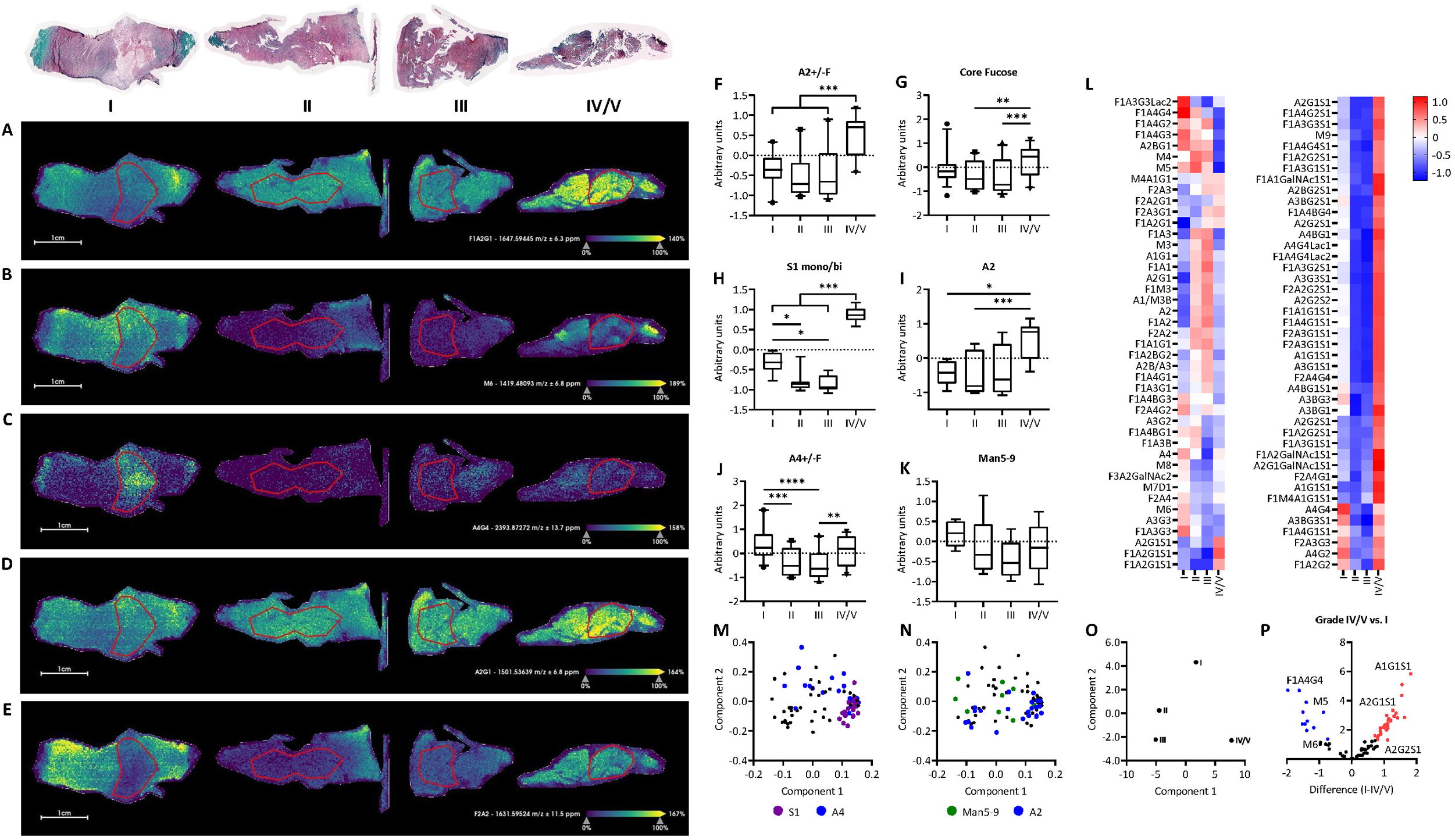
*N*-glycan imaging of the human IVD in degeneration. A-E) Glycan ion maps of representative glycan traits. Region of interest, nucleus pulposus, marked by red outline. F-K) Individual glycans were grouped into the following glycosylation traits; biantennary +/− outer arm fucosylation (A2+/−F) (F), monosialylated, core fucosylated glycans (G), mono/biantennary glycans (S1 mono/bi)(H), biantennary excluding outer arm fucosylated species (A2)(I), tetraantennary +/− outer arm fucosylation (A4+/−F) (J), higher oligomannose - from Man5 to Man9 (Man5-9)(K). Two-way ANOVA, Tukey’s post hoc test, *p <0.05, ***p <0.001. L) Z-score normalised heatmap of glycan abundance across grades. M-P) Principal component analysis separating grades of IVD degeneration and loadings highlighting the contribution of H) sialylated and tetraantennary glycans and I) Man5-9 and biantennary glycans towards each condition. J) Heatmap of z-score normalised glycans across all grades of degeneration.

### Exploratory Analysis of the IVD Proteome

Interpreting changes in *N*-glycosylation requires characterisation of the proteome from which the *N*-glycans were cleaved, to account for non-enzymatic changes in glycosylation secondary to protein synthesis and degradation. Protein expression of the human IVD in degeneration was analysed by LC-MS/MS. Healthy tissue digests contained more proteins than degenerated tissues. A total of 1,590 proteins were identified across all samples. 427 of the 1,590 were verified across three or more samples per group. The relative expression of all proteins quantified was normalised to total protein content with intensity derived from the sum of all corresponding peptide intensities. A PERL script was employed to calculate theoretically observable peptides through *in silico* digestion to normalise protein intensities. Inclusion parameters for all tryptic peptides were set at 6-30 amino acids, while missed cleavages were excluded. The full list of all proteins with significantly altered expression is summarised in Table S4. The most abundantly expressed proteins consist of ECM constituents including glycoproteins, proteoglycans and collagens (Fig. 6B). In NP degeneration, 16 proteins (Top 5: IL17B, CLEC3A, MATN3, VIT, MPO) were significantly downregulated while 28 proteins were upregulated (Top 5: VTN, APOA4, C1QC, POSTN, APCS) vs healthy NP (Fig 6E). In AF degeneration, 11 proteins were significantly downregulated (Top 5: AOC, HAPLN3, CLEC3A, THBS4, IL17B) while 14 proteins were upregulated (Top 5: PLA2G2A, APOA4, APCS, TTR, TIMP3) vs healthy AF (Fig 6F).

**Figure 6.**
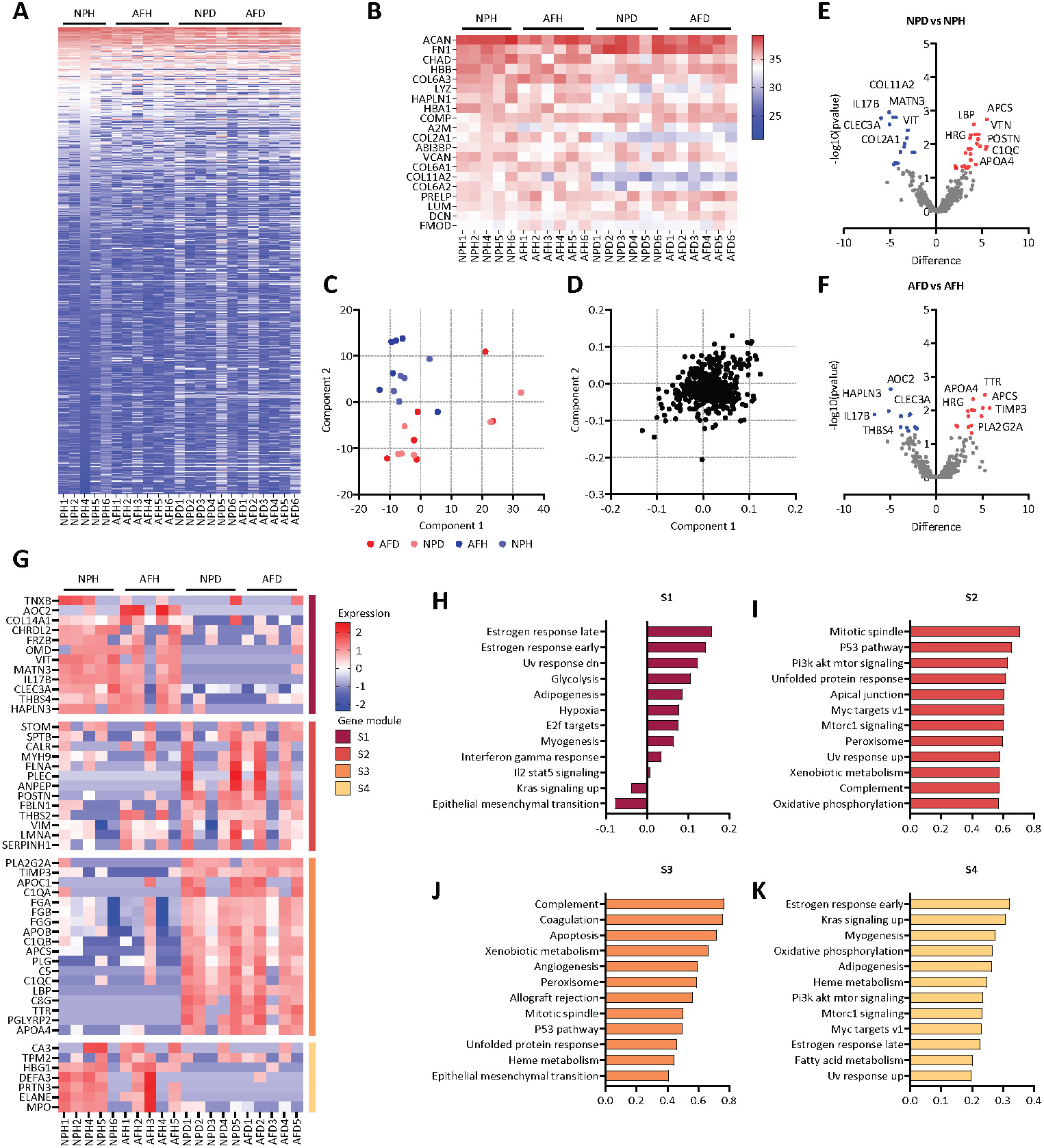
IVD degeneration is associated with acute phase signalling and apoptosis. A) Heatmap of protein’s relative abundance across IVD samples and (B) top 20 most highly expressed proteins. C+D) Principal component analysis and loadings separating healthy and degenerated IVD samples. (E+F) Volcano plots of significantly differentially expressed proteins across the NP (E) and AF (F) in degeneration, two-sided t-test, FDR <0.05. G) Subunit separation based on experimental groups. H-K) Most highly activated pathways in each subunit.

PCA was performed to gain a global overview of the proteomic data on the complete set of filtered proteins. The first two principal components described a combined variance of 47.3%, with PC1 and PC2 contributing 29.7% and 17.6%, respectively (Fig. 6C-D). Clear demarcation was observed between healthy NP, healthy AF, and degenerated tissues. PC1 provides a strong separation of healthy and degenerated tissues.

Pathway analysis of the degenerated AF revealed an upregulation of complement activation (GO:0030449) and downregulation of collagen fibril organisation (GO:0030199), collagen biosynthesis and modifying enzymes (R-HSA-1650814). The degenerated NP revealed upregulation of complement cascade (R-HSA-977606), RAF/MAP kinase cascade (R-HSA-5673001), complement activation (GO:0030449), regulation of ERK1 and ERK2 cascade (GO:0070372); with subsequent downregulation of collagen biosynthesis and modifying enzymes (R-HSA-1650814), assembly of collagen fibrils and other multimeric structures (R-HSA-2022090), and skeletal system development (GO:0001501).

The Clustered Heatmap (Fig 6G) is a 2-way unsupervised hierarchical clustering technique that clusters the expression matrix along rows and columns, clustering similar genes and samples. The functional annotation of clusters demonstrates Hallmark collection pathway regulation for each gene module (Fig 6H-K). Gene modules S2 and S3 are associated with a degenerated proteome and have significant pathway annotation scores for the p53 pathway, PI3K Akt signalling, unfolded protein response and apoptosis. Gene modules S1 and S4 are associated with a healthy proteome and have positive annotation scores for oestrogen response, KRAS signalling, hypoxia and myogenesis.

### RNA Sequencing Revealed Inflammatory Pathway Mediation in Sialylation Inhibited Nucleus Pulposus Cells

To characterise the cellular response to sialylation inhibition in IVD degeneration, an *in vitro* model using human nucleus pulposus cells was established as previously described (12). Healthy cells (H-CON) were stimulated with inflammatory cytokines (IL-1β, TNF-α and IL-6) to establish a pro-inflammatory response (H-CYTKN). The treatment group (H-TREAT) included the co-incubation of 3Fax-peracetyl Neu5Ac (Neu5Ac-inhib) in the pro-inflammatory culture conditions. In parallel, degenerated cells (D-CON) were also treated with Neu5Ac-inhib (D-TREAT), where no cytokines were used in this culture as it is presumed that these cells are conditioned to a chronic pro-inflammatory state.

An overview of the sample groups is provided through a principal component analysis (PCA) (Fig.7A). In the PCA plot, PC1 accounts for 42.41% of the variation while PC2 accounts for 10.54%. Both components separate groups based on cytokine stimulation only. Few DEGs were identified when comparing degenerated and healthy NP cells (D-CON vs H-CON; up 646, down – 715) (Fig 7B), while 1575 DEGs were confirmed in degenerated cells treated with Neu5Ac-inhib (D-TREAT vs D-CON); up – 712, down – 863) (Fig 7C) (Table S5). A total of 5174 differentially expressed genes (DEGs) were identified in healthy NP cells under cytokine stimulation (H-CYTKN vs H-CON; up – 2709, down - 2465) (Figure 7D). Similarly, 2881 DEGs were identified when inflamed cells were treated with Neu5Ac-inhib (H-TREAT vs H-CYTKN; up – 1151, down – 1730) (Fig 7E). The pro-inflammatory genes upregulated in cytokine-stimulated cells, decreased their expression with Neu5Ac treatment (Fig 7F-G).

**Figure 7.**
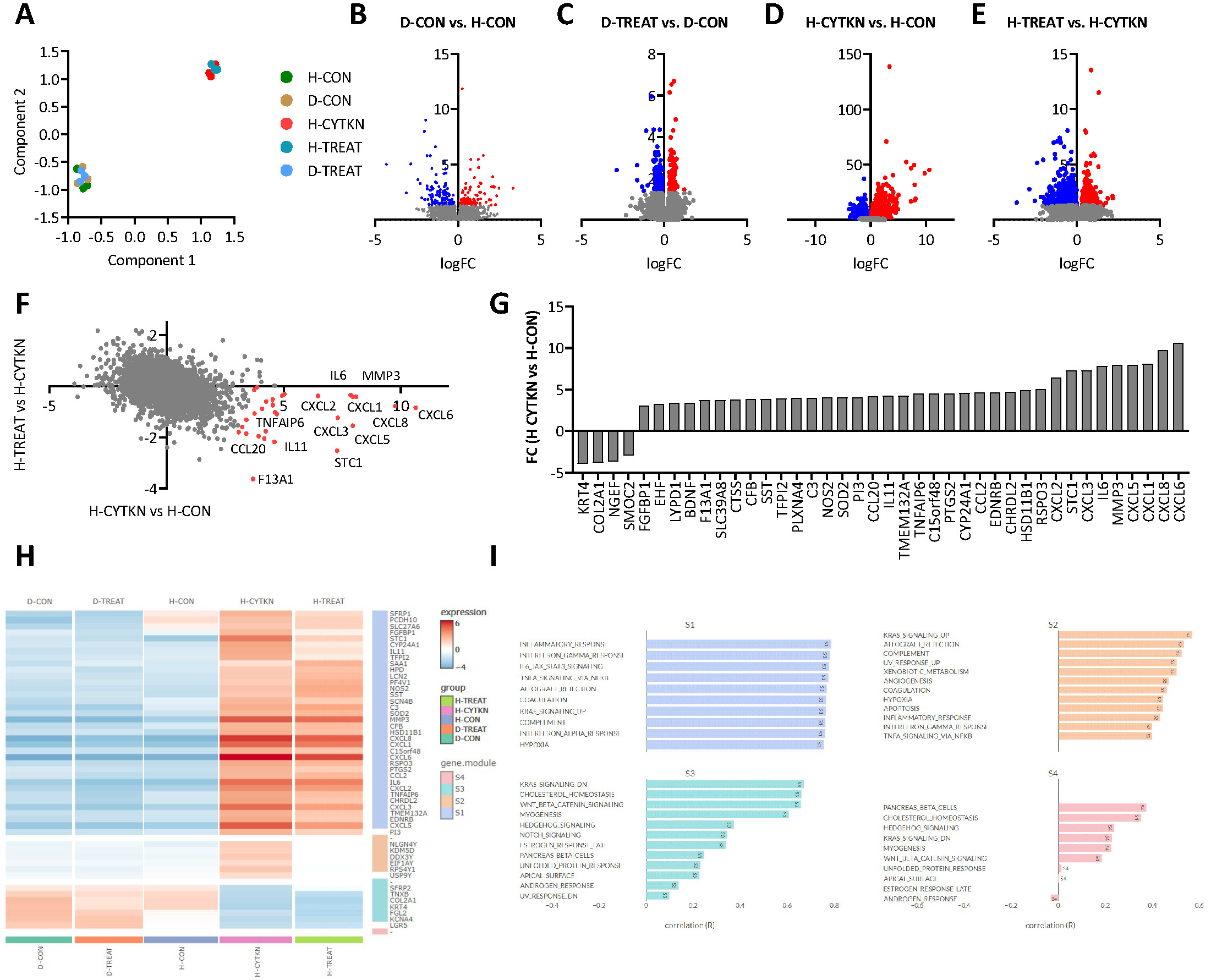
Transcriptional data from human NP cells under inflammatory and glycosylation inhibitor treated conditions. A) PCA analysis of five conditions: H-CON (healthy NP cells), D-CON (degenerated NP cells), H-CYTKN (healthy NP cells +cytokine stimulation), H-TREAT (healthy NP cells +cytokine stimulation + 3F-peracetyl Neu5Ac), and D-TREAT (degenerated NP cells + 3F-peracetyl Neu5Ac). B-E) differential gene expression across experimental conditions; Upregulated genes are highlighted in red, and downregulated genes are highlighted in blue. F-G) Differential expression analysis comparing common genes dysregulated under cytokine stimulation and Neu5Ac-inhib. H-I) Grouped subset analysis with associated pathway analysis using unsupervised hierarchal clustering.

Pathways such as Inflammatory response, IL6JAK STAT, TNF-α and complement activation are enriched in upregulated subunit clusters in cytokine stimulation (Fig 7I). There was a significant decrease in TNF-α signalling, inflammatory response and epithelial-mesenchymal transition in Neu5AC-inhib treated cells (Fig.8). The same response to sialylation inhibition is not observed in non-stimulated degenerated cells. There is an upregulation of glycosyltransferases in cytokine-stimulated cells vs healthy NP cells (Fig 9A-B), however, this upregulation is not observed in degenerated NP cells (Fig. 9C-D). This is likely due to the resolution of cellular stress in culture conditions vs *in vivo* conditions.

**Figure 8.**
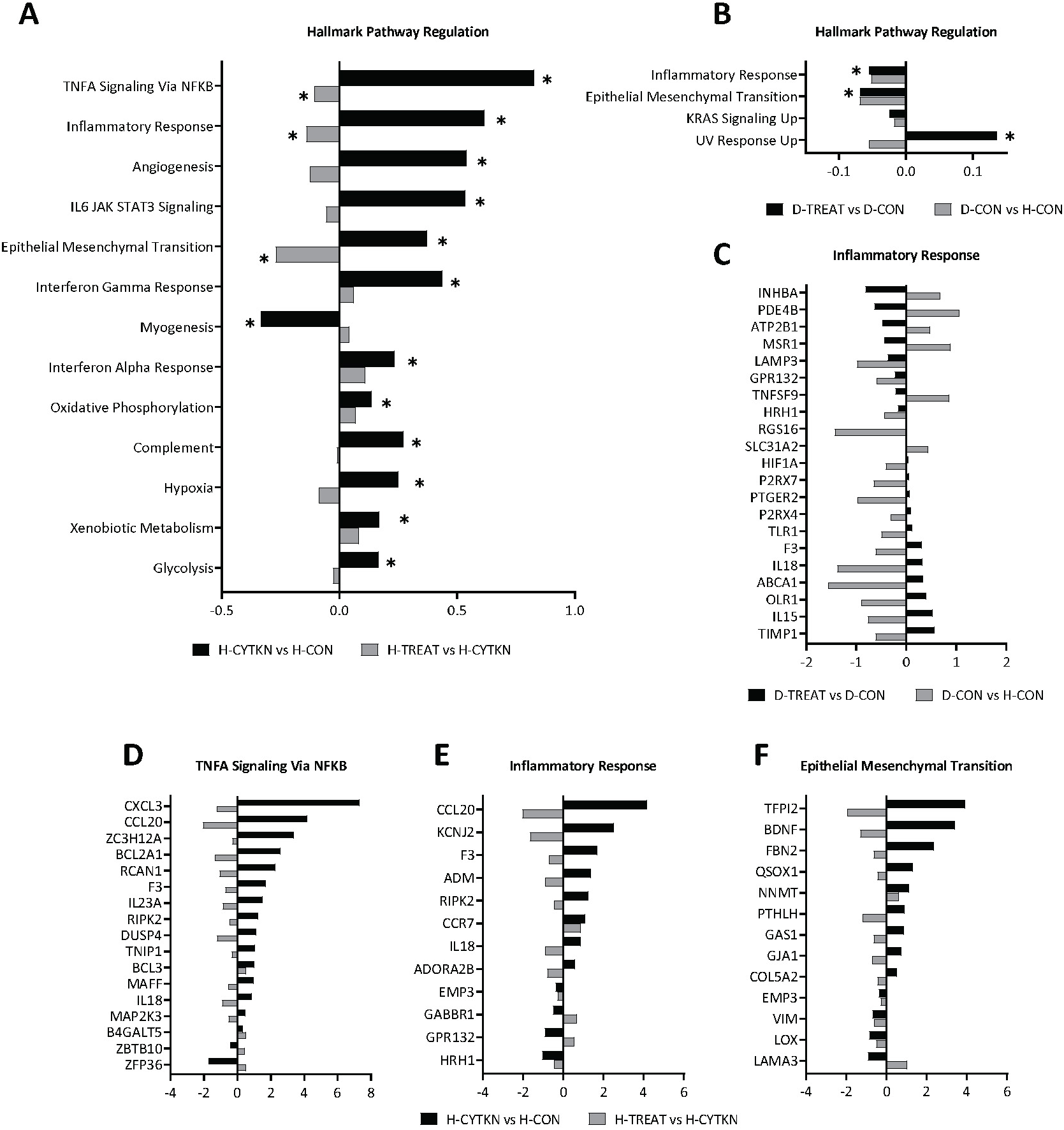
Transcriptional data from human NP cells under inflammatory and glycosylation inhibitor treated conditions. A+B) Hallmark pathway regulation demonstrating significantly dysregulated pathways in cytokine and cytokine/Neu5Ac-inhib conditions (H-TREAT vs H-CYTKN and H-CYTKN vs H-CON) (A) and unstimulated/Neu5Ac-inhib treated cells (D-TREAT vs D-CON and D-CON vs H-CON) (B), false discovery rates (FDR) <0.2. C) Significantly downregulated genes in the inflammatory response in Neu5Ac-inhib treated degenerated NP cells, p <0.05. D-F) Significantly dysregulated genes in TNFA signalling (D), Inflammatory response (E) and Epithelial Mesenchymal Transition (F) in Neu5Ac-inhib treated cytokine-stimulated NP cells, p <0.05.

**Figure 9.**
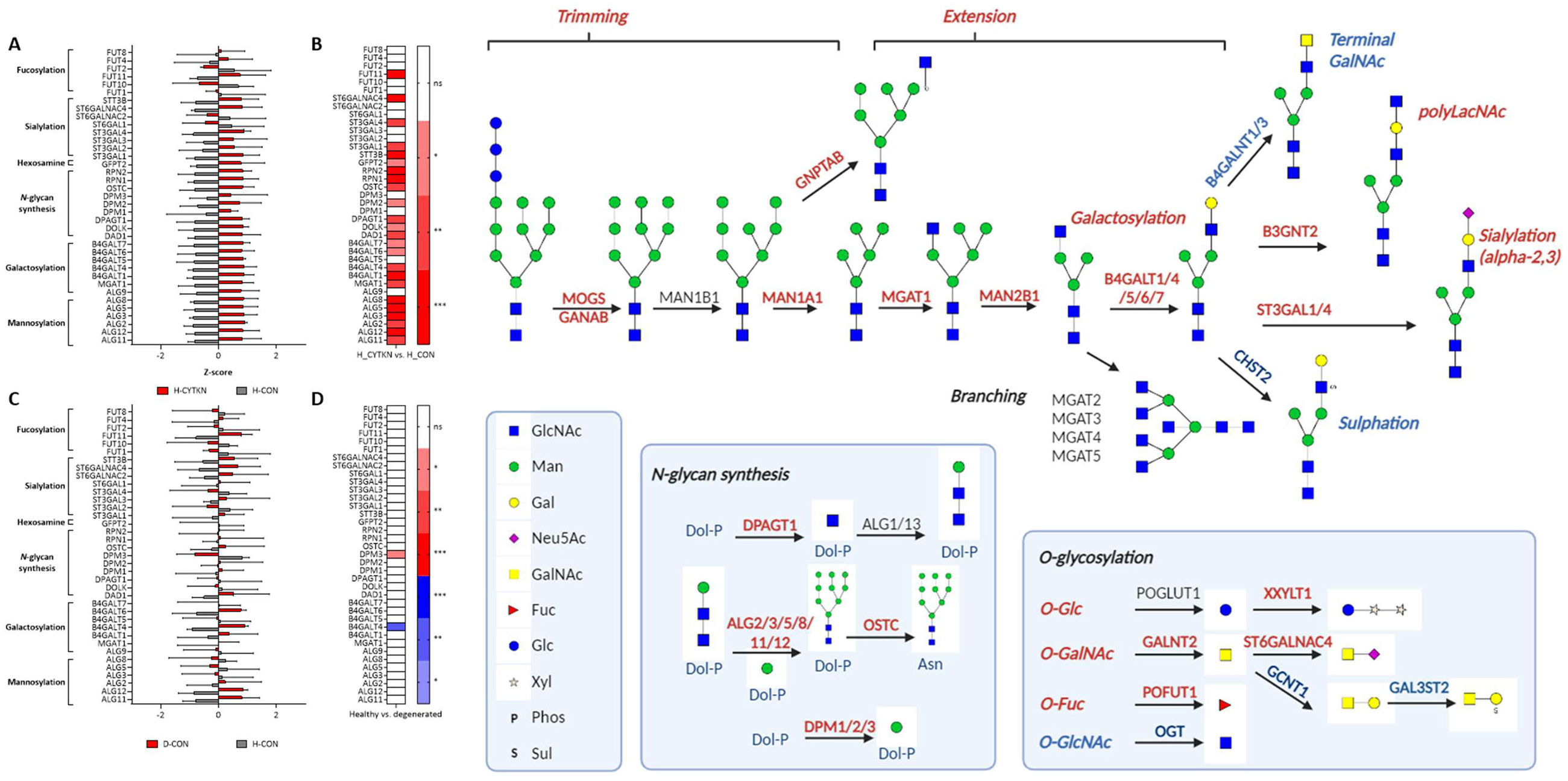
Overview of glycosylhydrolases and glycosyltransferases in *N*-glycan synthesis and decoration under cytokine stimulation in human NP cells. A+B) Glycosyltransferase expression in cytokine stimulation, significance interval presented in the heatmap, *p <0.05, **p <0.01, ***p <0.001 (B). C+D) Glycosyltransferase expression in degeneration, significance interval presented in heatmap, **p <0.01, ***p <0.001 (D). Upregulated (red) and downregulated (blue) genes are highlighted, two-sided t-test, FDR <0.05.

**Figure 10.**
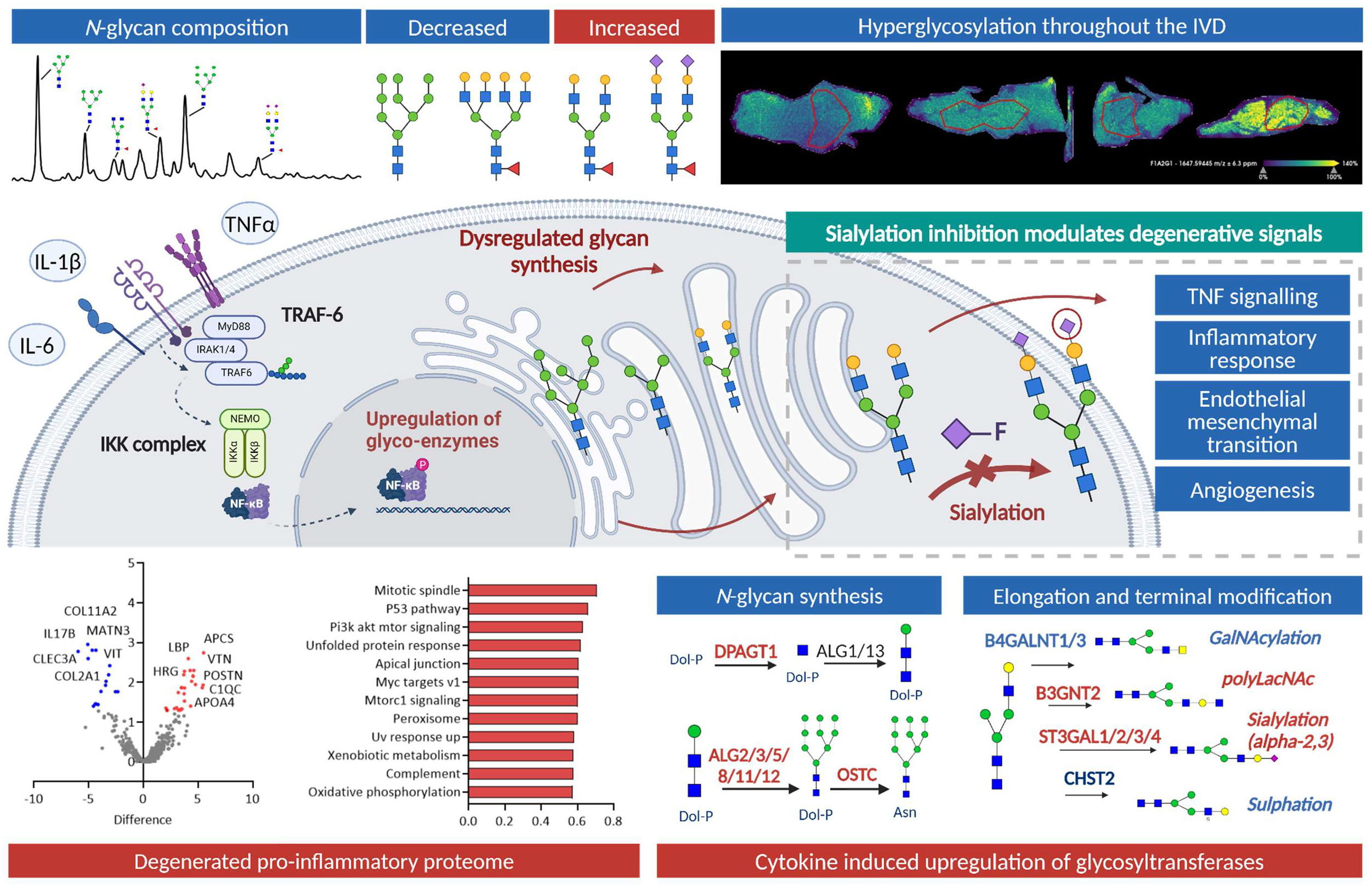
Glycosylation in IVD degeneration; cause and effect of aberrant glycan synthesis. Overview of glycomic and proteomic analysis in the context of IVD pathophysiology. The changes in glycosylation towards mono/bi-antennary sialylated glycans are associated with a pro-inflammatory proteome. Inhibitor studies have demonstrated an inhibition in TNF and inflammatory response pathways in a hypo-sialylated state.

## Discussion

Extensive cellular glycosylation analyses of the human intervertebral disc (IVD) have been limited in previous research efforts. However, it is increasingly recognised that glycosylation plays a vital role in various biological processes. Developmental morphological changes are often correlated with alterations in carbohydrate expression on cell surfaces and in the extracellular matrix (ECM). Studies in both rat and human models have suggested that disruptions in normal glycosylation patterns may contribute to developmental malformations (5,17,18). Despite this, a notable gap exists in understanding glycosylation patterns during human vertebral morphogenesis, maturity and disease (19).

Sialylation, specifically α-(2,6) sialylation through ST6GAL1 is required for somatic cell reprogramming and its downregulation is associated with decreased reprogramming efficiency (20). Therefore, the observed upregulation of sialylation and fucosylation within the IVD may signify phenotypic changes in resident cells. Similarly, the upregulation of outer arm fucosylation has been indicated in the functional regulation of TIMPs. The ability of TIMP-1 to inhibit MMPs is at least in part regulated by the outer arm fucosylation of its *N*-glycans (21). While TIMP1 expression increases in the degenerated disc, as shown here and previously described (22), it is plausible that its MMP-binding capacity is compromised due to increased outer arm fucosylation.

It is crucial to note that the *N*-glycosylation site function is unique to each glycoprotein sequence. This uniqueness, coupled with regulating the Golgi *N*-glycan-branching pathway, determines surface glycoprotein levels (23). Notably, this pathway is highly sensitive to intracellular hexosamine flux, particularly in the production of tri- and tetra-antennary *N*-glycans. These complex glycans have been shown to bind to galectins and form a molecular lattice that hinders glycoprotein endocytosis, thereby regulating surface protein expression (23).

In recent years, there has been growing interest in developing glycosylation inhibitors as potential therapeutic agents (24). Within the context of intervertebral disc degeneration (IVD), understanding the potential of glycosylation inhibitors is of particular relevance. In this study, we sought to comprehensively characterise the *N*-glycome of the human intervertebral disc (IVD) with spatial and temporal resolution. Our primary goal was to investigate the dysregulation of glycans in the context of IVD degeneration. To achieve this, we employed a multi-faceted approach that combined lectin histochemical analysis, glycan profiling, and proteomic analysis.

The results of initial lectin histochemical analysis in this study were also consistent with previous glycosylation profiling in bovine (25) and murine (9) IVD injury models and ovine IVD ageing (11). Differential regulation of glycosylation detected by lectins spurred the first *N*-glycan characterisation of the human IVD in degeneration. Increased expression of sialylation and fucosylation on UPLC-MS aligns with the observations made with lectin histochemistry (SNA; α-(2,6)-sialylation, UEA; α-(1,2)-linked fucose) reflecting the increased expression of sialyltransferases/fucosyltransferases (ST3GAL1/4, ST6GALNAC4, FUT11) in cytokine-induced inflammation (Fig.9), supported by previous studies in inflammation in fibrous tissues (14,26). The upregulation of sialyltransferases (ST6GAL1) and expression of α-(2,3) and α-(2,6)-sialylated glycosylation motifs have been characterised in human cartilage degeneration, as indicated by the binding of several lectins (27). MALDI-IMS analysis validated the UPLC results in age-matched comparisons. Proteomic analysis revealed an upregulation in acute phase response signalling which has been characterised previously to upregulate the expression of sialyltransferases and fucosyltransferases in connective tissues (28). These proteomic changes highlight a complex interplay between protein expression and glycosylation in IVD degeneration. The observed alterations in ECM constituents and glycoproteins may contribute to the glycosylation changes identified in the study.

*In vitro*, we explored how Neu5Ac-inhib affects glycosylation inhibition through different gene set enrichment analyses of transcriptomic data. RNA sequencing analysis in the given experimental groups revealed activation of inflammatory signalling in cytokine-stimulated NP cells. Co-incubation of NP cells with Neu5Ac-inhib altered the expression of signalling pathways associated with ECM organisation, cell-cell interactions, adheren junction, *N*-glycan synthesis and processing and various GO term enriched pathways identified through pathway analysis. The resultant hyperglycosylation states identified in this model are likely due to the upregulation in glycan synthesis and turnover, yet specific sialyltransferases were also significantly upregulated. ST3GAL1, found to be upregulated with cytokine stimulation, has previously been shown to be upregulated in an *in vitro* model of osteoarthritis (29). This can, in combination with ST6GAL1, modulate Galectin-1 reactivity.

Hallmark pathways such as TNF signalling, angiogenesis, IL6 JAK STAT3 signalling, and interferon-gamma response are enriched in cytokine-stimulated NP cells. The observed changes in gene expression patterns, particularly the downregulation of proinflammatory genes in response to Neu5Ac-inhib, indicate the potential of glycosylation modulation to mitigate inflammation, a significant factor in IDD pathogenesis.

Studies have shown that nutrient flow, driven by highly active growth-promoting receptors, can stimulate cell arrest and differentiation pathways. This effect is mediated by increased presentation of sparsely decorated *N*-glycoproteins with mono- and biantennary glycosylation (30). Notably, we observed a similar phenomenon in the human intervertebral disc (IVD), where a decrease in branching was evident. This suggests a close metabolic link between cellular growth regulation and the number and branching of *N*-glycans. The degree of *N*-glycan branching appears to serve as a molecular mechanism responsive to hexosamine flux, thereby modulating the metabolic transition between cellular growth and arrest. Importantly, this regulation seems to be influenced more by hexosamine flux and nutrient availability, particularly considering the reduced nutrient supply in degenerated IVDs, rather than being solely mediated by glycosyltransferase activity. Furthermore, the prevalence of oligomannose-type structures in healthy tissue may indicate cellular pluripotency, as such structures are recognised as markers of pluripotent stem cells (31,32). It is plausible that the greater abundance of oligomannose motifs in healthy tissue results from endoplasmic reticulum (ER) expansion during translational processes, rather than being indicative of ER stress (33).

## Conclusion

In conclusion, our comprehensive investigation into glycosylation patterns in intervertebral disc degeneration (IVD) sheds light on the critical role of glycosylation in this complex process. We observed distinct alterations in glycan expression, notably hypersialylation and decreased branching, which are associated with IVD degeneration. These changes indicate phenotypic shifts in resident IVD cells and have implications for regulating inflammatory responses and extracellular matrix dynamics. Furthermore, the underlying altered proteomic expression provides several indications towards the interplay between proteomic and glycomic homeostasis in IVD degeneration. Overall, transcriptomic analysis of cytokine stimulation in NP cells elucidates the functional role of sialylation in inflammatory signalling. Our findings highlight the importance of glycosylation in IVD biology and suggest that targeting specific glycosylation motifs, such as sialylation, may hold therapeutic potential in mitigating inflammation and oxidative stress associated with IVD degeneration. Overall, this research advances our knowledge of the glycomic mechanisms underlying IVD degeneration and opens new avenues for developing glyco-functionalised therapies to restore disc physiology.

## Materials and Methods

### Material and Reagents

AcroPrep™ Advance 96-filter plates and 10-kDa MWCO microcentrifuge filtration devices were purchased from Pall® Life Sciences, USA. Protogel™ was purchased from National Diagnostics™, France. 0.45 µm Millex-LH filters and C18 ziptips were purchased from Merck™, USA. 1 mL tuberculin BD Plastipak^©^ precision syringes were purchased from Medguard®, Ireland. Polypropylene 2 mL deep 96-well blocks and PhyNexus™ C18 phytip® columns were purchased from Fisher Scientific™, USA. Silverseal™ aluminium was purchased from Greiner Bio-One®, Austria. Plate seals were purchased from Cruinn Diagnostics®, Dublin. Sealing Mats were purchased from Phenomenex®, USA. Radio immunoprecipitation assay (RIPA) buffer, optimal cutting temperature (OCT) compound embedding medium and Superfrost plus slides were purchased from Thermo Fisher Scientific®, USA. Ammonium hydroxide solution was purchased from Honeywell Fluka™, USA. Leucine encephalin standard was purchased from Waters®, USA. All lectins were purchased from Vectors labs®, USA. PNGase F (P0709L) was purchased from New England Biolabs®, USA. All exoglycosidases were purchased from either New England Biolabs® or Prozyme®, USA. 3Fax-Peracetyl Neu5Ac was purchased from Merck. Human recombinant IL-1β, IL-6 and TNF-α were purchased from Peprotech. Primocin was purchased from InvivoGen. All other reagents were purchased from Merck®, USA, unless otherwise specified.

### Collection and Processing of Human Tissues

Human tissues were collected from three hospitals in Galway: Galway University Hospital, Merlin Park Hospital and Bon Secours Hospital, and from Our Lady’s Children’s Hospital, Crumlin, with ethical approval from NUI Galway Ethics Committee (CA 269), Health Service Executive and Crumlin Research Ethics Committee. These tissues were used for UPLC N-glycan analysis, lectin microarray analysis and proteomic analysis. Healthy human NP cells for transcriptomic analysis were isolated from discs procured during spinal realignment surgery for adolescent idiopathic scoliosis (donor age range: 9-16 years old, Pfirrmann grade I). Degenerated cells were isolated from discs during discectomy and fusion surgery (donor age range: 36-72 years old, Pfirrmann grade IV/V). All participants signed consent forms before tissue collection. In instances where the patient was under the age of 16, a consent form was signed by the patient’s parent or guardian. In cases where the child was between 16 and 18, the child signed an assent form and a consent form from a parent/guardian. Degenerated intervertebral discs were harvested from patients with diagnosed intervertebral disc degeneration. These discs were procured during discectomy and microdiscectomy procedures. Healthy intervertebral discs were procured from adolescent patients undergoing spinal realignment surgery. Microdiscectomy is often performed in the lumbar disc levels along with facetectomy to increase lumbar mobility before pedicle screw fixation. The AF and NP were identified by the surgeon and segregated into different sample cups, wrapped in damp gauze and stored at 4°C until collection within four hours. For MALDI analysis, age matched controls for Grade I-III were obtained from the Netherlands. Collection of intervertebral disc specimens adhered to medical ethical regulations set out by University Medical Centre Utrecht (Utrecht, The Netherlands).

Tissues were further processed in a tissue culture hood. Samples were washed three times with Hank’s balanced salt solution (HBSS) for three minutes each wash (to remove residual blood/plasma, which would interfere with glycan analysis). 100 mg of tissue was transferred to 1 mL RIPA buffer with protease inhibitor cocktail (1%), stored at 4°C for one hour, and then frozen at −80°C. 100 mg was flash-frozen, and the remaining tissue was used for cell isolation.

Once all samples were collected for *N*-glycan analysis, the RIPA stored samples were thawed at 4°C. A stainless steel bead was added to each Eppendorf® and the tubes were added to the Qiagen Tissuelyser LT™ set at 50 Hz, for 80 minutes. If tissue was not completely homogenised after 80 min, homogenisation was continued for further 30 minutes. Once homogenisation was complete, Eppendorf tubes were spun at 16000 g for 20 minutes at 4°C. The supernatant was separated from the pellet and transferred to LoBind™ Eppendorf tube. The supernatant was dried in a vacuum centrifuge (Savant™ SPD131DDA SpeedVac™ Concentrator, ThermoFisher®) and stored at −80°C. This soluble fraction of tissue homogenate was further processed for *N*-glycan isolation.

### Classification of Disc Degeneration

The Thompson grading system is used to assess IVD degeneration, based on NP morphology, AF and end plate intactness, and osteophyte formation. Thompson grading was employed for samples received from The Netherlands for the preliminary lectin histochemical-based pilot study, where MRI images were not available to assess IVD degeneration using Pfirrmann grading. Sagittal T2-weighted magnetic resonance (MR) images of the lumbar spine were used to grade morphologic disc degeneration based on MR signal intensity, disc structure, the distinction between AF and NP, and disc height. The Pfirrmann grading system was employed to assess the extent of disc degeneration according to the following table, where grade varies from I to V. This grading system was used for samples collected in Ireland that were subsequently used for *N*-glycan analysis.

### Lectin Microarray Analysis

The standard procedure used for the glycoprotein labelling: 50 µg by isolated glycoprotein of each sample (prior to gel block isolation in *N*-glycan analysis) was added to 10 µL of 10X phosphate buffer (500 mM, pH 8.3) and diluted to 99 µL with PBS. Afterwards, the solution was treated with 0.3 µL of Alexa-647 NHS ester (10 µg/µL in DMSO) for one hour at room temperature. The excess dye was quenched by the addition of 0.7 µL of 1 M Tris buffer. The fluorescently labelled glycoprotein solutions were not purified and were directly used in the lectin array analysis. Printed slides stored at −20°C were retrieved and quenched by immersion in a 50 mM ethanolamine solution in borate buffer (50 mM, pH 8.5) for 45 minutes at room temperature, and the quenched surface was then passivated by incubation in PBS containing 0.5% Tween-20, 0.4 mg/mL BSA, 1 mM CaCl_2_, 1 mM MgCl_2_ and 1 mM MnCl_2_ for 45 minutes at room temperature. The slide was dried by centrifugation. The glycoprotein samples were added to the corresponding wells and the incubation was carried out for 3.0 h at r.t. Samples were then aspirated, and the slide was washed with PBS for 5 minutes and dried by centrifugation and scanned. Microarray data interpretation: The images obtained from the G265BA microarray scanner (Agilent Technologies®) were analysed with Pro Scan Array Express software (PerkinElmer®) to determine fluorescence intensities. For each subarray, all fluorescence values were normalised to the highest single fluorescent value before combining lectin replicates to a single value. From six printed lectin spots per subarray, the maximum and minimum values were removed to generate an average from the median four values. Values from duplicate analyses for each sample were combined for the average and standard deviation values.

### *N*-Glycan Isolation – In Gel Block

Gels were made around the dried tissue homogenate to immobilise the *N*-glycosylated proteins made up of 64.7% Protogel, 32.30% Gel buffer (1.5M TRIS pH8.8) and 3% of 10% sodium dodecyl sulphate (SDS) solution. Gel volume was added until the homogenate was completely dissolved (100 μL for the sample). Ammonium peroxisulphate (APS) and *N,N,N,N’*-Tetramethyl-ethylenediamine (TEMED) were added to the gel solution at a ratio of 1:35. Gels were allowed to set for 20 minutes. They were then transferred to the freezer for 10 minutes for ease of cutting in the next step. Gels were chopped into 1 mm^3^ pieces on a clean glass plate using a clean scalpel. Gels were washed with 20 mM sodium bicarbonate solution and acetonitrile to wash impurities and unreacted gel components. The gels were reduced and alkylated using dithiothreitol (DTT, 0.5M) and iodoacetamide (IAA, 100 mM), respectively. This step removed disulphide bonds across peptides to expose *N*-glycan linkages to the peptide to be cleaved by PNGase F. The gel was further washed through dehydration and rehydration washes and PNGase F (1,250 units/mL) was added to each gel and incubated at 37°C overnight. Next, the glycans were eluted from the gels through several sonication steps and washes. The elution was filtered using a 0.45 μm LH Millipore filter and a 1 mL syringe and dried overnight in the vacuum centrifuge. Dried glycans were reduced in formic acid for 40 minutes, resuspended in 2-aminobenzamide (2AB) solution and incubated at 65°C for 30 minutes for labelling through reductive amination (34). The solution was then transferred to Whatman 3MM chromatography paper and excess 2AB was washed away with acetonitrile (35). The glycans were eluted with water and dried in the vacuum centrifuge.

### Analysis of Human *N*-glycome on Hydrophilic Interaction Liquid Chromatography/ Ultra Performance Liquid Chromatography (HILIC-UPLC)

UPLC was performed using a BEH Glycan column (1.7 μm particles in 2.1×150 mm, Waters) on an Acquity UPLC equipped with a temperature control module and an Acquity fluorescence detector. Solvents A and B were composed of 50 mM formic acid adjusted to 4.4 pH with ammonia solution and acetonitrile (Sigma-Aldrich Acetonitrile E CHROMASOLV for HPLC, far UV), respectively. The column was maintained at a temperature of 40°C. Solvent A was applied using a linear gradient from 30-47% over 30 minutes followed by 47-70% A and finally 30% A to complete each run (35). Samples were prepared in 70% acetonitrile. The excitation wavelength was set at 330 nm with detection at 420 nm. Calibration was performed using hydrolysed and 2AB-labelled glucose oligomers as an external standard creating a dextran ladder that was used for every run (36).

### Weak Anion Exchange (WAX) – UPLC Determination of Sialylation

WAX – UPLC was completed using a Waters Acquity UPLC separations module complete with an Acquity HPLC fluorescence detector through the Empower Chromatography Workstation. The analytical column used was a Waters DEAE anion exchange column (75 x 7.5 mm, 10 µm particle size). Mobile phase A consisted of 20% v/v acetonitrile in water, (Milli-Q water, quality > 18.2 MΩ, TOC content < 5 ppb). Mobile phase B consisted of 0.1 M ammonium acetate buffer pH 7.0 in 20% v/v acetonitrile. A linear gradient of 0 to 5% solvent A over 12 minutes at a flow rate of 1 mL/min was applied, followed by 5−21% solvent A over 13 minutes and then 2−50% A over 25 minutes, 80−100% A over 5 minutes followed by five minutes at 100% (36). The standard used for charged state chromatogram annotation is 10% v/v fetuin N. Normal human serum (NHS) *N*-glycans labelled were used as an additional standard to ensure the instrument is correctly calibrated. Fluorescence detection was set at excitation/ emission wavelengths of *λ*_ex_ = 330 nm and *λ*_em_ = 420 nm, respectively.

### Exoglycosidase Digestions

Exoglycosidases were procured from Prozyme (San Leandro, CA) or New England Biolabs (AMF, BKF, GUH) (Hitchin, Herts, U.K.). The isolated 2AB-labelled glycans were digested for 18 hours at 37°C in 10 µL 50 mM sodium acetate buffer, pH 5.5 (except jack bean α-mannosidase (JBM) digestion which requires 100 mM sodium acetate, 2 mM Zn^2+^, pH 5.0). The following enzymes were used: almond meal α-fucosidase (AMF, EC 3.2.1.111), 40 mU/mL; *Arthrobacter ureafaciens* sialidase (ABS, EC 3.2.1.18), 0.5 U/mL; bovine kidney α-fucosidase (BKF, EC 3.2.1.51), 800 U/mL; bovine testes β-galactosidase (BTG, EC 3.2.1.23), 1 U/mL; β-*N*-acetylglucosaminidase cloned from *S. pneumonia*, expressed in *Escherichia coli* (GUH, EC 3.2.1.30), 8 U/mL (Prozyme) – 400 U/mL (NEB); Jack bean hexosaminidase (JBH, 3.2.1.52), 10 U/mL; Jack bean mannosidase (JBM, EC 3.2.1.24), 60 U/ mL; *Streptococcus pneumoniae* sialidase (NAN1, EC 3.2.1.18), 5 U/mL; *Streptococcus pneumoniae* β-galactosidase (SPG, EC 3.2.1.23), 0.4 U/mL. Glycosidases were removed after incubation by filtration through 10 kDa protein-binding EZ™ filters (Millipore Corporation). *N*-glycans were then analysed by UPLC as previously described (36).

### Liquid Chromatography-Mass Spectrometry-Fluorescence of *N*-glycans

Glycan profiles were obtained by negative ion nanoelectrospray LC-MS, performed by Acquity® UPLC system through a BEH Glycan Column (150 x 1.0 mm i.d., 1.7 µm particles), coupled to a Waters® Xevo® G2 QTOF system. The data acquisition was performed with the instrument set as previously described with augmentation as outlined in Table 2.1 (37). Data acquisition and analysis were performed using MassLynx™ (Waters®, Milford, MA, USA). The FLD excitation/emission spectra were set to 320 nm and 420 nm, respectively. The sample injection was 8 µL (75% MeCN). The flow rate was set to 0.15 mL/min and the column temperature was maintained at 60 °C. A linear gradient was applied as follows: 0.0 min 28% A 72% B, 1.0 min 28% A 72% B, 31.0 min 43% A 57% B, 32.0 min 45% A 55% B, 36.0 min 28% A 72% B, 40.0 min 28% A 72% B.

### Sectioning of IVD Tissue for Histological Examination

Discs from healthy and degenerated tissue were obtained from Utrecht University in the Netherlands. The IVD specimens were collected according to the medical ethical regulations (protocol 12-364) of the University Medical Centre Utrecht (Utrecht, The Netherlands). Isolated discs were fixed in 10% formalin for 48 hours, and decalcification of the vertebral bones was performed using Kristensen’s decalcifying solution (18% (v/v) acetic acid/3.5% (w/v) sodium formate) at 4°C. Following decalcification, samples were then washed in running tap water for 12 hours and transferred to 20% (w/v) sucrose solution until submerged at 4°C. Optimal cutting temperature (OCT) compound was used to embed tissues, which were then snap-frozen in an isopentane bath with liquid nitrogen. Samples were stored at −80°C until sectioning at 10 µm on a cryostat (Leica CM1850).

### *N*-glycan MALDI-FTIMS/FTICR

Formalin-fixed paraffin-embedded (FFPE) tissues were sectioned at 5 µm and mounted onto charged slides (Superfrost™ Plus). Slides were subsequently dewaxed and rehydrated before undergoing antigen retrieval using citraconic anhydride buffer (25 µL citraconic anhydride, 2 µL HCl and 50 mL HPLC grade water, pH. 3.0-3.5) as previously described [19]. After antigen retrieval, slides were desiccated and underwent digestion by recombinant PNGase F enzyme (Serva Electrophoresis GmbH, Heidelberg, Germany) applied using a TMSprayer™ (HTX Technologies LLC., Chapel Hill, NC). Enzyme solution was sprayed onto the slides at 25 µL/min for 15 passes at 45°C. Slides were incubated in a humidified chamber for two hours at 37°C. After incubation, slides were desiccated and 7 mg/mL CCA matrix was applied by the TMSprayer™ at 100 µL/min for 10 passes at 80°C. Slides were stored in a desiccator overnight until further analysis by MALDI-FTICR MS. Released *N-*glycan ions were detected using a MALDI FTICR scimaX™ (Bruker Daltonics, Germany) for accurate, high resolution mass analysis, operating in positive ion mode with a Smart Beam II laser operating at 1000 Hz and a laser spot size of 20 µm. The signal was collected at a raster width of 200 µm between spots. A total of 10 laser shots were collected to form each pixel, calibrated to release a total of 5× 10^8^ ions from each spot. Following the acquisition, data was processed and images of expressed glycans were generated using SCiLS™ Lab 2024 software (Bruker Daltonics, Germany), where ions in the range of 650-3400 m/z were analysed and statistically evaluated the expression across sample cohorts normalised to the total ion count. Observed mass/ charge ratios were searched against glycan databases using GlycoWorkBench (38). Represented glycan structures were generated in GlycoWorkBench and composition was determined by accurate mass (<10 ppm error), and previous structural characterisation elucidated by UPLC for the human *N-*glycan profile described above. SCiLS 2018 (Bruker Daltonics) imaging software was used to further analyse glycan expression and statistically evaluate expression across sample cohorts normalised to the total ion current.

### Histopathological overview

After MALDI analysis, the matrix was removed from the slides by dipping in 100% acetone until slides were clear. Slides were then washed in demineralised water for five minutes before being subjected to Weigert’s Haematoxylin for five minutes. Slides were then washed in running tap water for 10 minutes, rinsed in distilled water and counterstained with filtered 0.4% Fast Green for four minutes, subjected to 1% acetic acid two times for a total of five minutes and stained with 0.125% aqueous Safranin for five minutes. The sections were then dehydrated in 96% Ethanol for one minute twice, followed by 100% ethanol for five minutes and xylene two times for five minutes and mounted using DPX mountant.

### Proteomic Evaluation of Human IVD

After *N*-glycans had been released during glycan isolation, the gels were rehydrated, washed and dried in a vacuum centrifuge. 200 µL of trypsin in 50 mM ammonium bicarbonate (ratio enzyme: substrate 1:50) was added to each sample and incubated at 37°C overnight. The digested peptides were eluted with 1% formic acid 50% AcN: H_2_O through multiple washes and sonication steps. Trifluoroacetic acid (TFA) was added to the sample to a final concentration of 0.1% before purification with a C18 Zip tip. Samples were dried to approx. 10-20 µL volume and frozen at −20°C until LC-MS analysis. Samples were run on a Q Exactive™ Hybrid Quadrupole-Orbitrap™ Mass Spectrometer (UCD Conway, Dublin). Briefly, samples were dissolved in 0.1% formic acid, loaded onto a fused silica emitter (75 µm ø), pulled with a laser puller and packed with reverse-phase media. An increasing acetonitrile gradient was applied over 47 minutes at a 250 nL/min flow rate. The instrument was operated in positive-ion mode with a potential of 2,300 V applied to the frit and a capillary temperature of 320°C. A high-resolution (70,000) MS scan across 300–1,600 m/z was performed with Q Exactive to identify the eight most intense ions, followed by MS/MS analysis with higher-energy collisional dissociation. Protein identification was performed by searching the raw data against the *Homo sapiens* (Human) subset of the UniProt Swiss-Prot database (UP000005640_9606.fasta) using MaxQuant computational platform (Max-Planck-institute of Biochemistry). Label-free quantification was performed based on specified peptides with specified enzymatic cleavage for Trypsin/P with fixed modification of carboxymethylation and deamidation, as outlined previously (39). Each peptide used for protein quantification was subject to FDR filtering of <1% to be accepted for analysis. MaxQuant data was exported to Perseus® software for proteomic analysis (40). The output of differentially expressed proteins was further investigated using Ingenuity Pathway Analysis® (IPA; Qiagen®, Redwood City, USA). Data was also independently analysed using PEAKS studio (Bioinformatics Solutions® Inc.) for peptide identification and label-free quantification to validate MaxQuant identification and quantification.

### Transcriptomics

Degenerated cells (discectomy patients, Pfirrmann grade V) and healthy cells (adolescent idiopathic scoliosis, Pfirrmann grade I) were procured intraoperatively with consent. AF and NP were identified and separated in the theatre by the surgeon. The tissue was weighed based upon an addition to a pre-weighed falcon tube containing PBS (pH 7.4) and 1 % P/S. After washing, the tissue was minced and a 0.2 % Pronase solution (pH 7.4) was added for 1 h at 37 °C on a shaker plate at 300 rpm. The tissue was washed twice with PBS and incubated in collagenase solution (100 U/mL collagenase in α-MEM and 10 % FBS) (pH 7.4). The sample was digested overnight and filtered through a 100 µm cell strainer. The cell solution was centrifuged and washed before cell counting. Cells were plated at a density of 10,000 cells/cm^2^. Cells were cultured in a complete medium containing α-MEM, 10 % FBS and 1 % P/S in hypoxia (1% oxygen, 5% CO_2_) at 37 C. All experiments were performed on cells within passage 4 or sooner due to the loss of cell phenotype in 2D culture conditions. Healthy Control (H-CON: NP cells extracted from healthy IVD), Degenerated Control (D-CON: NP cells extracted from degenerated IVD), Healthy Cytokine (H-CYTKN: healthy NP cells stimulated with cytokine cocktail – IL1β, IL-6, TNFα), Healthy Treated (H-TREAT: combination of healthy NP cells, cytokine cocktail and Neu5Ac-inhib), Degenerated Treated (D-TREAT: degenerated NP cells and Neu5Ac-inhib). (41)). The cytokine-containing medium was refreshed at 48 h. The ‘Degenerated Treated’ group does not contain the cytokine cocktail, as these cells were hypothesised to be preconditioned having already been exposed to an inflammatory microenvironment in vivo.

### RNA Sequencing

#### RAW Data / Quality Control

The compressed paired-end human mRNA-seq data in fastq format over 80 gigabytes from Illumina PE150. The ‘fastqc’ v.3.2 quality control tool (Babraham Bioinformatics) was applied. Each group consisted of three biological replicates (five experimental groups), and each biological replicate was divided into two paired-end files. The “multiqc” tool v.2.1.4 generated duplication reads, average GC content, and entire sequences but also sequence quality histograms, per sequence quality scores, per sequence, quality content and adapter content graphs (42). ‘trim-galore v.4.2’ performed adapter trimming and low-quality read filtering. Moreover, ‘fastp’ tools v.2.1 were used for filtering low-quality cells (43). After completing these steps, each replicate has over 99 % of the read-passing filter. Alignment, assembly and quantification were performed using Kallisto” tools v.2.3. (ENSEMBL Human cDNA file (the transcriptFASTAfile: Homo_sapiens.GRCh38). We applied the “tximport” package to import transcript-level abundances from Kallisto quantification tools into R studio and convert into gene counts for downstream analysis. The biomart “ensembl” database and the “hsapiens_gene_ensembl” dataset were used for connecting and mapping transcript IDs to gene IDs. “DESeq2” v.1.36.0 (44) was used to perform differential expression analysis. We explored four different experiment data analyses between two groups: H-CYTKN vs H-CON, D-CON vs H-CON, H-TREAT vs H-CYTKN and D-TREAT vs D-CON. A PCA plot that identifies the samples as clusters or patterns and relationships in the data was generated through the PCAtools R package. In the differential gene analysis, the adjusted p-value cutoff (FDR) and alpha value were 0.05, the Benjamini-Hochberg method was applied for adjusting p-values, and independent hypothesis weighting was chosen as the performance of independent filtering through IHW v.1.24.0 Bioconductor package (45). A shrinkage method called “Adaptive Permutation based Empirical Bayes Gene-wise Linear Models” known as ‘apeglm’ (46) were applied to account for unreliably large log fold change estimates. Biological and statistically significant differentially expressed upregulated and downregulated genes are determined by log_2_FC >1 and adjusted *p*-value <0.05. Different gene set enrichment in GO, KEGG and Reactome pathways was analysed using “fgsea” v.1.24.0 R package with default parameters (normalised enrichment score - <2.2 />-2.2, FDR 0.05). The results are described in dot plot and the heatmap shows the differentially expressed genes in each specific enriched gene set. Finally, the over-representation methods of “clusterprofiler package” v.4.8.1 (47) were utilised to identify functionally enriched pathways with parameters of 0.05 adjusted p-value and q-value cut-off and the Benjamini-Hochberg method as pAdjustmethod after changing gene-IDs into Entrez-IDs, producing dot plots and category plots.

#### Statistical Analysis

Graphs and figures were created using GraphPad Prism 8 Software. Statistical analyses were conducted as described in the figure legends using Prism 8 software.

## Acknowledgements

Mr Pat Kiely (CHI Crumlin, Ireland) and Prof. Laura Creemers (Utrecht University, Netherlands) for their assistance in the provision of human tissues.

## Author Contributions

Conceptualization: KJ, AP, Methodology: KJ, RD, MMD, RS, AP, Investigation: KJ, AS, AMP, Supervision: AD, MMD, RS, AP, Writing—original draft: KJ, AS, Writing—review & editing: KJ, AS, MMD, RS, AP.

## Competing interests

None.

**Data and Materials availability**

